# Female mice exhibit a more sensitive automated squint response to pain induced by CGRP and amylin

**DOI:** 10.1101/2021.05.26.445893

**Authors:** Brandon J. Rea, Levi P. Sowers, Abigail L. Davison, Aaron M. Fairbanks, Anne-Sophie Wattiez, Pieter Poolman, Randy H. Kardon, Andrew F. Russo

**Author notes:** corresponding author, 51 Newton Road, Iowa City, IA 52245. These authors contributed equally to this manuscript (Brandon J. Rea, Levi P. Sowers).

## Abstract

We developed an automated squint assay using both black C57BL/6J and white CD1 mice that measured the interpalpebral fissure area between the upper and lower eyelids as an objective quantification of pain. In C57BL/6J mice, we observed a squint response to increasing doses of a migraine trigger, the neuropeptide CGRP, including a significant response in female mice at a dose below detection by the manual grimace scale. Using the automated software, both C57BL/6J and CD1 mice lacked a detectable photic blink response. The CGRP-related peptide amylin induced squinting behavior in female mice, but not males. These data demonstrate that an automated squint assay can be used as an objective, real-time continuous-scale measure of pain that provides higher precision and real-time analysis compared to manual grimace assessments.

## INTRODUCTION

Pain assessment in humans and animals is critical for the development and monitoring of new pain therapeutics. While pain is inherently subjective to the individual experiencing it, current assessment techniques rely on verbal responses and questionnaires. Pain assessment can be even more problematic for individuals that lack the ability to communicate effectively, such as children and patients with neurodevelopmental disorders [3; 11]. The difficulty in translating pain assessment tools to the bench and back to humans extends to limitations on the use of mice as an experimental platform for understanding pain. To overcome these shortcomings, researchers have sought continuous objective measures of pain through the analysis of tissue biomarkers, electrical potentials, imaging procedures and assessment of muscle tension [6; 19]. These approaches are rational, but often lack correlation with other signs of pain and recording of these components in a clinical setting is often difficult. An exciting advancement in rodent pain assessment is the recent development of high-speed photographic monitoring of paw withdrawal touch-based assays, which has revealed time-resolved features previously missed by observer monitoring [9]. However, translation of a paw withdrawal assay to patients is not feasible. The measurement of dynamic changes of facial features activated during a grimace response has been recognized as an alternative, viable objective biomarker for pain that can be observed across species, including humans [14].

Manual scoring of facial features to quantify grimace in laboratory animals is effective but is time intensive, requires training of graders to reduce subjectivity, and is limited to a noncontinuous three-level scale of severity (0, 1, 2) [12; 16]. Despite the challenges of facial grimace scoring, the assay has proven quite useful and highly translatable to humans [19; 22]. To address these challenges, video-based capture of facial images have been manually scored [12] or scored by trained convoluted neural networks for the presence or absence of facial pain [20]. However, many of these platforms rely on methods that can only detect extremes in facial pain, making it difficult to identify small changes in pain severity.

In this study, we sought to validate an automated video-based squint assay to measure pain by leveraging our previous finding that orbital tightening, or squint, is the principal component of the mouse grimace score [16]. The automated squint assay was validated by comparing to manual measurements of grimace and orbital tightening in response to the neuropeptide CGRP, which is known to cause squinting and other components of grimace in mice [16]. We then used the automated squint software to assess the photic blink response in mice, with validation of automated results using electromyography (EMG). Finally, the automated assay was used to test the sex-dependent effect of the CGRP-related peptide amylin, which binds one of the two CGRP receptors [8] and the amylin analog pramlintide and amylin have recently been shown to induce migraine in patients and migraine-like symptoms in mice, respectively [5].

## MATERIALS AND METHODS

### Animals

Wildtype C57BL/6J black (https://www.jax.org/strain/000664) and CD1 white mice (http://www.criver.com/products-services/basic-research/find-a-model/cd-1-mouse) were used for development of the automated squint model. In the CGRP dose response squint assay, cohorts of 20 C57BL/6J mice were used for each CGRP dose (10 males, 10 females) and vehicle (11 males, 9 females). C57BL/6J (10 males, 4 females) and CD1 mice (10 males, 9 females) were used for the automated photic blink assay. Three nestin/hRAMP1 double transgenic mice and one littermate control were used in the recording of orbicularis oculi EMG in response to light and air puff [24]. In the amylin squint assay, amylin treated C57BL/6J (11 males, 10 females) and vehicle (8 males, 8 females) cohorts were used. Mice were 10-14 weeks old, with an average weight of 26 g for C57BL6/J mice and 30 g for CD1 mice. All strains of mice were housed in a temperature-controlled vivarium on a 12-hour light cycle with food and water ad libitum. C57BL/6J mice were housed in groups of 5, CD1 mice were housed in groups of 4, and the chronically implanted EMG electrode nestin/hRAMP1 double transgenic and control mice were single housed. All behavioral experiments were performed between 8:00 A.M. and 5:00 P.M. after a minimum of 1-week habituation in the animal facility. All procedures followed the ARRIVE guidelines and were approved by the University of Iowa Institutional Animal Care and Use Committee and implemented in accordance with the standards set by the National Institutes of Health.

### Intraperitoneal drug administration

Rat α-CGRP (Sigma-Aldrich, St. Louis, MO) was diluted in Dulbecco phosphate buffered solution (PBS, HyClone) and administered via intraperitoneal (IP) injection with a 30 g x 0.5-inch needle in the following quantities: 0.01 mg/kg, 0.05 mg/kg, 0.1 mg/kg, or 10 mL/kg PBS alone as a vehicle. Rat amylin (Bachem, Torrance, CA) was diluted in Dulbecco PBS (HyClone) and administered via IP injection at 0.5 mg/kg or 10 mL/kg PBS alone as a vehicle. Mice were handled gently with no anesthesia during injections. Investigators were blinded to drug treatment with all injections performed by either B.J.R. or L.P.S. Mice injected with CGRP or vehicle were allowed to recover for 30 minutes in their home cage before testing. Mice injected with amylin or vehicle were allowed to recover for 15 minutes in their home cage before testing.

### Development of video image capture for mouse facial detection

Video recordings of C57BL/6J and CD1 mice sampled at 1 frame every 0.1 s (10 frames per second) were used for training the automated facial detection software. Mice were acclimated to a customized gentle collar restraint prior to experimentation as previously described [16] to fix camera distance in order to constrain video magnification changes as well as to decrease struggle and head movement. Acclimation sessions for C57BL/6J and CD1 mice were 20 minutes each for 3-4 or 4-5 sessions, respectively. Acclimation was individually determined per mouse by willingness to remain still for at least 10 minutes. Mice that did not acclimate to the custom restraint were excluded. Eight synchronized cameras were employed from varying vantage points for each recording session (IDS Imaging UI-3240ML-NIR, Obersulm, Germany) with infrared light to visualize the face and ensure landmark detection. A custom graphical software tool was used to manually locate and place a predetermined set of 20 facial anatomical landmarks on video frames displaying varying degrees of mouse expression. Following facial landmarking, a proprietary algorithmic approach (FaceX, LLC, Iowa City, IA) was used to derive these landmarks in newly recorded video frames and identify the face and eye. The software incorporated a combination of deep learning and shape regression models to detect, align, and label facial and eye landmarks. The software-reported accuracy of the landmark placement was made by a secondary shape regression model that estimated a tracking error rate as the root-mean squared error of the predicted landmarks. After direct comparison of software-reported tracking error and manual assessment of eye landmark alignment by inspecting each frame of forty 5-minute video recordings, individual frames containing a tracking error rate of >15% were excluded. This error rate was where the software tracking error aligned with all manually assessed video recordings.

### Automated measurement and analysis of squinting behavior, facial grimace, and photic blink

For automated squint analysis, the custom gentle collar restraint and acclimation protocol previously described was employed. Following a 5-minute baseline video recording and squint measurement in room light, CGRP (0.01 mg/kg, 0.05 mg/kg, 0.1 mg/kg) or vehicle (PBS, 10 mL/kg) was administered to C57BL/6J mice via IP injection and were returned to their home cage. Mice were restrained once more 30 minutes post-injection and recorded for squint assessment over 5 minutes in room light. Pixel area measurement for the right eye palpebral fissure was derived every 0.1 seconds (10 frames per second) in the recordings using the trained facial detection software with the resulting values compiled with custom MATLAB script. Individual frames containing a tracking error rate of >15% were excluded. In comparing manual grimace scoring versus automated squint assessment, the frame with largest pixel area delta between baseline and treatment in pixel area for each CGRP cohort was selected and scored using the Mouse Grimace Scale [12] by three blinded individuals. For automated squint analysis with amylin (0.5 mg/kg) and vehicle (PBS, 10 mL/kg), C57BL/6J mice underwent the same protocol as CGRP-treated mice but were restrained and recorded for squint assessment 15 minutes post-injection. In the photic blink assay, CD1 and C57BL/6J mice were acclimated and restrained in the same manner and recorded in a dark room (∼0 lux), then exposed to a white light stimulus (1250 lux, 5600K, 1 sec), with pixel area measurement for the right eye palpebral fissure derived every 0.1 seconds.

### Implantation of EMG telemeter and recording of orbicularis oculi muscle activity

Mice were acclimated to the gentle collar restraint prior to surgery. Under aseptic conditions using a stereotaxic unit (Kopf Instruments, Tujunga, CA) and a rodent warming pad (Stoelting, Wood Dale, IL), mice were anesthetized with isoflurane (5% induction with 1 L/min oxygen, 1-2% maintenance with 1 L/min oxygen) and subcutaneously implanted with a radiotelemeter (DSI, New Brighton, MN), with leads in the orbicularis oculi muscle of the left eye. Each mouse was housed individually and monitored during recovery. After a 24-hour recovery period, mice were gently restrained and recorded in a dark room (∼0 lux), then exposed to a light stimulus (745 lux, 455 nm). Following light exposure, mice were reacclimated to the dark (∼0 lux) for 15 seconds before a second light stimulus (745 lux, 455 nm). Following this, an ipsilateral corneal air puff was delivered by B.J.R. or A.M.F. using a glass pipette and rubber bulb (1 mL). Changes in EMG activity for the light only and light plus air puff sessions were collected at 2,000 Hz for 1 to 2 seconds using DSI Acquisition software (DSI, New Brighton, MN). We applied a bandpass filter to the EMG data from 30-600 Hz together with a band-stop (or notch) filter at 60 Hz and all its harmonics in the bandpass interval (120, 180, 240, 300, 360, etc.) to suppress powerline noise. Data is presented as the root mean square (RMS) of the EMG activity in 100 millisecond (ms) windows (every 0.1 seconds).

### Statistical analysis

All statistical analyses were performed using GraphPad Prism 9.1.0 based on data expressed as mean ± SEM. Statistical details are presented in Table 1. Differences in change from baseline to treatment with CGRP and vehicle were determined by 2-way repeated-measure ANOVA: treatment (3 different CGRP concentration groups, 1 vehicle group, 4 groups total) and condition (baseline and treatment) followed by Šídák’s multiple comparisons test to compare treatment effect with respective baselines. Statistical comparisons of automated squint from dark to light conditions in photic blink experiments were analyzed with paired 2-tailed *t*-tests. Differences in change from baseline to treatment with amylin and vehicle were determined by 2-way repeated-measure ANOVA: treatment (amylin, vehicle, 2 groups total) and condition (baseline and treatment) followed by Šídák’s multiple comparisons test to compare treatment effect with respective baselines. All experiments were repeated with different cohorts in at least two independent sessions. Data available upon request.

**Table 1:** Statistical Analysis.

| Figure # | Analysis | Statistics |
| --- | --- | --- |
| Figure 2B<br>top row<br>left panel | Two-way repeated measure ANOVA<br>Interaction factor<br>Treatment factor (4 treatment groups)<br>Condition factor (Baseline, Treatment) | $F_{(3,76)}=9.743, p<0.0001$<br>$F_{(3,76)}=4.756, p=0.0043$<br>$F_{(1,76)}=61.49, p<0.0001$ |
| Automated<br>Squint | Šídák's multiple comparison<br>- Baseline vs. Treatment for PBS group (n = 20)<br>- Baseline vs. Treatment for CGRP 0.01 mg/kg group (n = 20)<br>- Baseline vs. Treatment for CGRP 0.05 mg/kg group (n = 20)<br>- Baseline vs. Treatment for CGRP 0.1 mg/kg group (n = 20) | $p<0.9999$<br>$p=0.0387$<br>$p<0.0001$<br>$p<0.0001$ |
| Figure 2B<br>top row<br>middle panel | Two-way repeated measure ANOVA<br>Interaction factor<br>Treatment factor (4 treatment groups)<br>Condition factor (Baseline, Treatment) | $F_{(3,76)}=12.33, p<0.0001$<br>$F_{(3,76)}=1.225, p=0.3066$<br>$F_{(1,76)}=33.57, p<0.0001$ |
| Mouse<br>Grimace<br>Scale | Šídák's multiple comparison<br>- Baseline vs. Treatment for PBS group (n = 20)<br>- Baseline vs. Treatment for CGRP 0.01 mg/kg group (n = 20)<br>- Baseline vs. Treatment for CGRP 0.05 mg/kg group (n = 20)<br>- Baseline vs. Treatment for CGRP 0.1 mg/kg group (n = 20) | $p=0.4390$<br>$p=0.3196$<br>$p<0.0001$<br>$p<0.0001$ |
| Figure 2B<br>top row<br>right panel | Two-way repeated measure ANOVA<br>Interaction factor<br>Treatment factor (4 treatment groups)<br>Condition factor (Baseline, Treatment) | $F_{(3,76)}=12.70, p<0.0001$<br>$F_{(3,76)}=1.789, p=0.1564$<br>$F_{(1,76)}=36.34, p<0.0001$ |
| Manual<br>Orbital<br>Tightening | Šídák's multiple comparison<br>- Baseline vs. Treatment for PBS group (n = 20)<br>- Baseline vs. Treatment for CGRP 0.01 mg/kg group (n = 20)<br>- Baseline vs. Treatment for CGRP 0.05 mg/kg group (n = 20)<br>- Baseline vs. Treatment for CGRP 0.1 mg/kg group (n = 20) | $p=0.4030$<br>$p=0.1979$<br>$p<0.0001$<br>$p<0.0001$ |
| Figure 2B<br>middle row<br>left panel | Two-way repeated measure ANOVA<br>Interaction factor<br>Treatment factor (4 treatment groups)<br>Condition factor (Baseline, Treatment) | $F_{(3,37)}=12.70, p=0.0028$<br>$F_{(3,37)}=3.954, p=0.0153$<br>$F_{(1,37)}=17.55, p=0.0002$ |
| Male<br>Automated<br>Squint | Šídák's multiple comparison<br>- Baseline vs. Treatment for PBS group (n = 11)<br>- Baseline vs. Treatment for CGRP 0.01 mg/kg group (n = 10)<br>- Baseline vs. Treatment for CGRP 0.05 mg/kg group (n = 10)<br>- Baseline vs. Treatment for CGRP 0.1 mg/kg group (n = 10) | $p=0.9834$<br>$p=0.9338$<br>$p=0.0043$<br>$p=0.0003$ |
| Figure 2B<br>middle row<br>middle panel | Two-way repeated measure ANOVA<br>Interaction factor<br>Treatment factor (4 treatment groups)<br>Condition factor (Baseline, Treatment) | $F_{(3,37)}=10.06, p<0.0001$<br>$F_{(3,37)}=0.8479, p=0.4766$<br>$F_{(1,37)}=12.84, p=0.0010$ |
| Male<br>Mouse<br>Grimace<br>Scale | Šídák's multiple comparison<br>- Baseline vs. Treatment for PBS group (n = 11)<br>- Baseline vs. Treatment for CGRP 0.01 mg/kg group (n = 10)<br>- Baseline vs. Treatment for CGRP 0.05 mg/kg group (n = 10)<br>- Baseline vs. Treatment for CGRP 0.1 mg/kg group (n = 10) | $p=0.1775$<br>$p=0.9810$<br>$p=0.0032$<br>$p<0.0001$ |
| Figure 2B<br>middle row<br>right panel | Two-way repeated measure ANOVA<br>Interaction factor<br>Treatment factor (4 treatment groups)<br>Condition factor (Baseline, Treatment) | $F_{(3,37)}=11.12, p<0.0001$<br>$F_{(3,37)}=1.990, p=0.1323$<br>$F_{(1,37)}=13.73, p=0.0007$ |
| Male<br>Manual<br>Orbital<br>Tightening | Šídák's multiple comparison<br>- Baseline vs. Treatment for PBS group (n = 11)<br>- Baseline vs. Treatment for CGRP 0.01 mg/kg group (n = 10)<br>- Baseline vs. Treatment for CGRP 0.05 mg/kg group (n = 10)<br>- Baseline vs. Treatment for CGRP 0.1 mg/kg group (n = 10) | $p=0.2086$<br>$p=0.9998$<br>$p=0.0020$<br>$p<0.0001$ |
| Figure 2B<br>bottom row<br>left panel | Two-way repeated measure ANOVA<br>Interaction factor<br>Treatment factor (4 treatment groups)<br>Condition factor (Baseline, Treatment) | $F_{(3,35)}=4.325, p=0.0107$<br>$F_{(3,35)}=3.512, p=0.0251$<br>$F_{(1,35)}=50.14, p<0.0001$ |
| Female Automated Squint | Šídák's multiple comparison<br>- Baseline vs. Treatment for PBS group (n = 9)<br>- Baseline vs. Treatment for CGRP 0.01 mg/kg group (n = 10)<br>- Baseline vs. Treatment for CGRP 0.05 mg/kg group (n = 10)<br>- Baseline vs. Treatment for CGRP 0.1 mg/kg group (n = 10) | $p=0.9293$<br>$p=0.0131$<br>$p<0.0001$<br>$p<0.0001$ |
| Figure 2B bottom row middle panel | Two-way repeated measure ANOVA<br>Interaction factor<br>Treatment factor (4 treatment groups)<br>Condition factor (Baseline, Treatment) | $F_{(3,35)}=4.225, p=0.0119$<br>$F_{(3,35)}=1.321, p=0.2831$<br>$F_{(1,35)}=20.64, p<0.0001$ |
| Female Mouse Grimace Scale | Šídák's multiple comparison<br>- Baseline vs. Treatment for PBS group (n = 9)<br>- Baseline vs. Treatment for CGRP 0.01 mg/kg group (n = 10)<br>- Baseline vs. Treatment for CGRP 0.05 mg/kg group (n = 10)<br>- Baseline vs. Treatment for CGRP 0.1 mg/kg group (n = 10) | $p=0.9899$<br>$p=0.3331$<br>$p=0.0022$<br>$p=0.0009$ |
| Figure 2B bottom row right panel | Two-way repeated measure ANOVA<br>Interaction factor<br>Treatment factor (4 treatment groups)<br>Condition factor (Baseline, Treatment) | $F_{(3,35)}=3.861, p=0.0174$<br>$F_{(3,35)}=1.273, p=0.2986$<br>$F_{(1,35)}=22.94, p<0.0001$ |
| Female Manual Orbital Tightening | Šídák's multiple comparison<br>- Baseline vs. Treatment for PBS group (n = 9)<br>- Baseline vs. Treatment for CGRP 0.01 mg/kg group (n = 10)<br>- Baseline vs. Treatment for CGRP 0.05 mg/kg group (n = 10)<br>- Baseline vs. Treatment for CGRP 0.1 mg/kg group (n = 10) | $p=0.9913$<br>$p=0.0894$<br>$p=0.0009$<br>$p=0.0036$ |
| Figure 3A right panel<br>C57BL/6J Automated Squint | Paired 2-tailed t-test Dark vs. Light<br>$n = 14$<br>$df = 13$<br>$t = 1.982$ | $p=0.0690$ |
| Figure 3B right panel<br>CD1 Automated Squint | Paired 2-tailed t-test Dark vs. Light<br>$n = 19$<br>$df = 18$<br>$t = 0.7991$ | $p=0.4347$ |
| Figure 5B left panel<br>Automated Squint | Two-way repeated measure ANOVA<br>Interaction factor<br>Treatment factor (PBS, Amylin)<br>Condition factor (Baseline, Treatment)<br>Šídák's multiple comparison<br>- Baseline vs. Treatment for PBS group (n = 16)<br>- Baseline vs. Treatment for Amylin 0.5 mg/kg group (n = 21) | $F_{(1,35)}=2.133, p=0.1531$<br>$F_{(1,35)}=3.380, p=0.0745$<br>$F_{(1,35)}=7.149, p=0.0113$<br>$p=0.6706$<br>$p=0.0068$ |
| Figure 5B middle panel<br>Male Automated Squint | Two-way repeated measure ANOVA<br>Interaction factor<br>Treatment factor (PBS, Amylin)<br>Condition factor (Baseline, Treatment)<br>Šídák's multiple comparison<br>- Male Baseline vs. Treatment for PBS group (n = 8)<br>- Male Baseline vs. Treatment for Amylin 0.5 mg/kg group (n = 11) | $F_{(1,17)}=0.1064, p=0.7483$<br>$F_{(1,17)}=2.188, p=0.1574$<br>$F_{(1,17)}=2.208, p=0.1556$<br>$p=0.7045$<br>$p=0.3286$ |
| Figure 5B right panel<br>Female Automated Squint | Two-way repeated measure ANOVA<br>Interaction factor<br>Treatment factor (PBS, Amylin)<br>Condition factor (Baseline, Treatment)<br>Šídák's multiple comparison<br>- Female Baseline vs. Treatment for PBS group (n = 8)<br>- Female Baseline vs. Treatment for Amylin 0.5 mg/kg group (n = 10) | $F_{(1,16)}=3.448, p=0.0818$<br>$F_{(1,16)}=4.253, p=0.0558$<br>$F_{(1,16)}=5.626, p=0.0306$<br>$p=0.9294$<br>$p=0.0118$ |

## RESULTS

### Automated squint tracking and analysis

The trained FaceX software algorithm was able to automatically detect the C57BL/6J face using the 20 facial landmarks (Fig. 1A, red dots, not all visible) from the 8 synchronized cameras (Fig. 1B). After detection of the face, the software automatically aligned and labelled the facial and eye landmarks and applied the final tracing to the eyelid borders (Fig. 1A, B, green line). By comparing pixel area values with time-stamped videos, we could identify any changes in pixel area resulting from struggle, movement, blinks, and eye opening (Fig. 1C, black arrows). The same analyses could be performed in white CD1 mice (Fig. 1D-F). Over 1,000 images were trained for the CD1 (white) mouse model and over 2,200 images for the C57BL/6J (black) mouse model. The C57BL/6J mice required more training images due to the difficulty of discerning a black eyelid border and palpebral fissure against black fur. The 8 camera angles pictured in Fig. 1B, E illustrate the angles chosen for facial landmarking. In this restraint model, camera 2 provided the best angle and position for detection (Fig. 1B, E, red box) and analysis (Fig. 1C, F) of squint and was used for all experiments. These data demonstrated we could accurately track facial features of both black and white mice.

**Figure 1:**
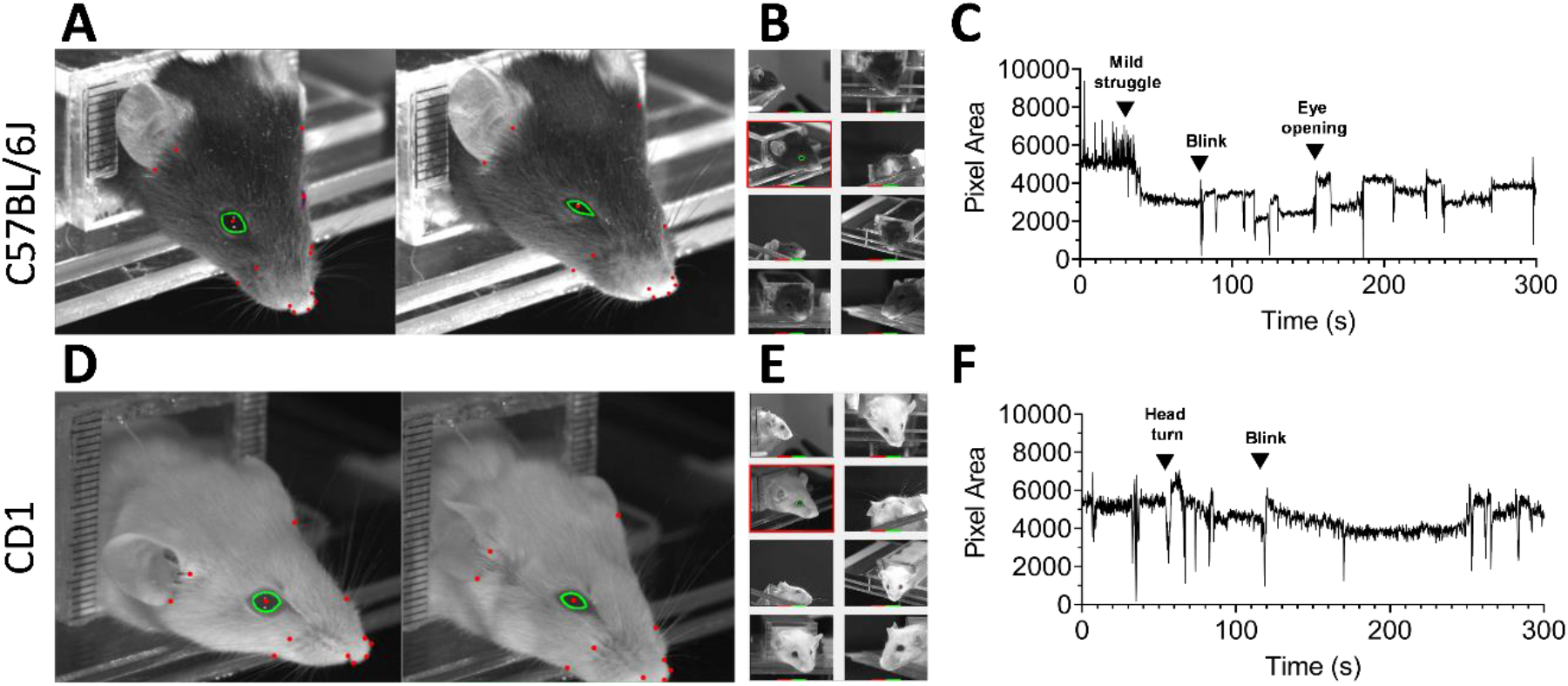
Framework and collection of automated squint data in C57BL/6J (black) and CD1 (white) mice. (**A**) C57BL/6J mice were held with a gentle collar restraint to allow denotation of facial landmarks (red dots, not all visible) at varying levels of facial pain presentation. The right frame shows automatically derived fissure narrowing (outlined by green border) in response to CGRP (**B**) The face was located using object detection software based on the facial landmarks from eight synchronized cameras to ensure the optimum pose was captured for automated squint analysis. A decision forest of binary features along with additional trained regressors located the eye and applied the final optimal tracing of palpebral fissure shown as green borders. When utilizing the gentle collar restraint, camera 2 (red box) provided the best angle to accurately track the right eye and measure pixel area of the palpebral fissure. (**C**) Pixel area of the right eye palpebral fissure was sampled every 0.1 seconds (10 frames per second) with notable events such as struggle, blink, head turn, and sustained eye opening denoted by black arrows. (**D-F**) Same as panels A-C except with CD1 mice.

### Automated squint versus manual orbital tightening and grimace scoring of CGRP-treated C57BL/6J mice

To validate the automated software, we assessed the squint and grimace responses to CGRP. We have previously reported that CGRP at 0.1 mg/kg causes a grimace and squint response in mice [16] but a dose-effect of grimace responses had not been performed. Analyzed with the automated software, CGRP-induced squint could be visualized over time (Fig. 2A). CGRP-administered mice displayed a significant squint with even the lowest dose of 0.01 mg/kg CGRP (Fig. 2B, top row, left panel). When the manual Mouse Grimace Scale was used to score mice at the point with the largest delta in pixel area between baseline and treatment in the automated assay (Fig. 2A black arrows), we confirmed a grimace at the higher CGRP doses, but not at the lowest dose (0.01 mg/kg CGRP) (Fig. 2B, top row, middle panel). When the grimace orbital tightening unit was analyzed separately from grimace, it did not show a significant difference at 0.01 mg/kg CGRP (Fig. 2B, top row, right panel).

**Figure 2:**
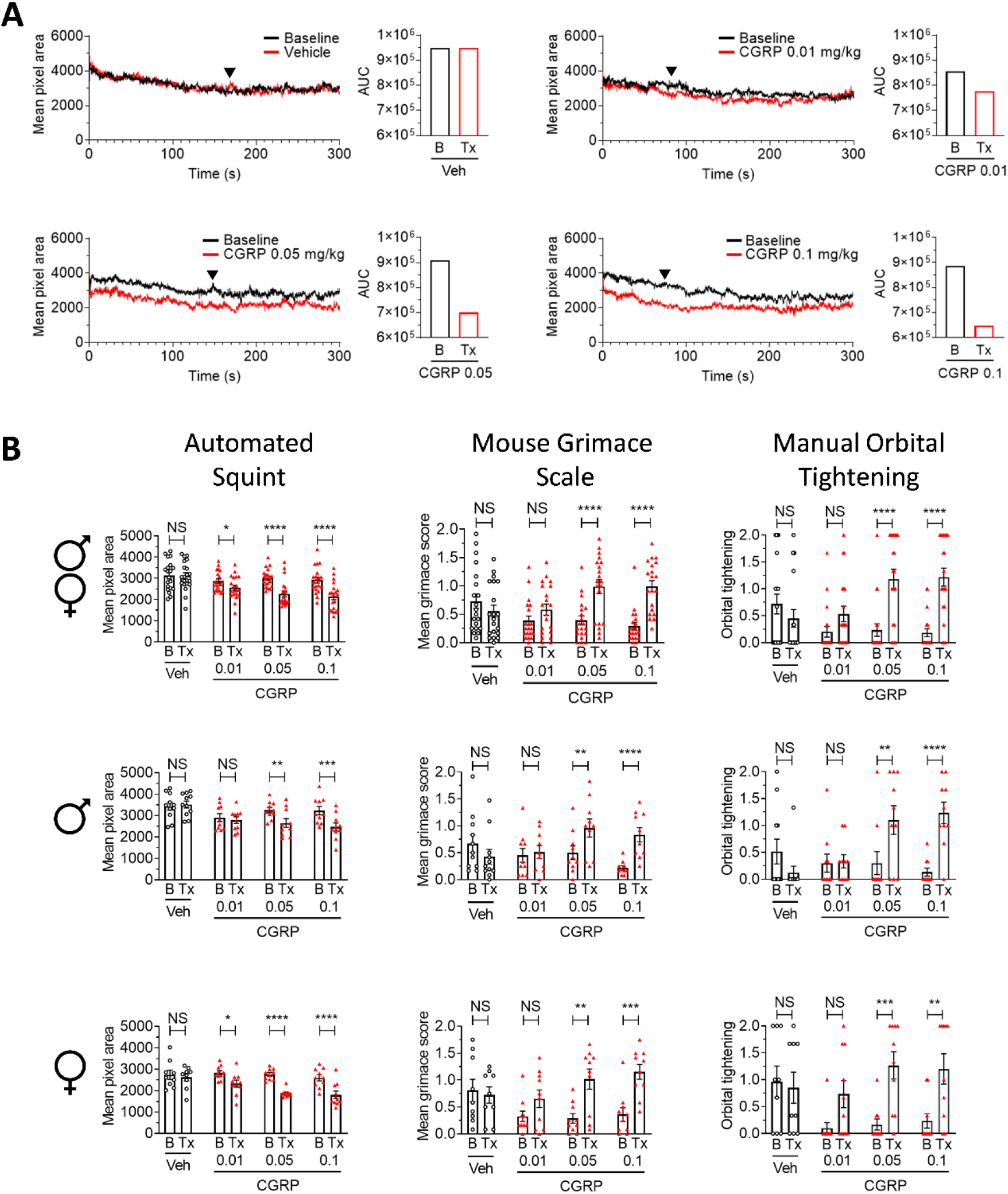
Automated squint analysis detects pain behavior with subthreshold levels of CGRP that Mouse Grimace Scale does not in C57BL/6J mice. (**A**) Mean pixel area over time for each baseline (5-minute recording, no injection, n = 20 for each) and respective treatment (5-minute recording, 30 minutes post injection) for all mice injected with either Veh (PBS, n = 20) or CGRP (0.01 mg/kg, 0.05 mg/kg, or 0.1 mg/kg, n = 20 for each). Black arrows indicate time synchronized frames with the greatest squint area delta between baseline and treatment independently assessed by three blinded individuals to maximize the likelihood of detecting a difference in Mouse Grimace Scale analysis. Area under the curve for the averages are shown as box plots to the right of the line graphs. (**B**) Mean overall pixel area, mean grimace scores, and mean orbital tightening action unit scores from the grimace analysis from panel A for all mice during baseline (B) and treatment (Tx) conditions. The data are shown for all mice (top row), male only (middle row, n = 11, PBS; n = 10 for each CGRP concentration) and female only (bottom row, n = 9, PBS; n = 10 for each CGRP concentration). Error bars indicate ± SEM. Two way repeated-measures ANOVA followed by Šídák’s multiple-comparison test to compare baseline and treatment conditions, *p < 0.05, **p < 0.005, ***p < 0.001, ****p < 0.0001.

Interestingly, when analyzed by sex, only females, not males, showed a significant automated squint response to 0.01 mg/kg CGRP (Fig. 2B, middle and bottom rows, left panels). Both sexes responded to the 0.05 mg/kg and 0.1 mg/kg CGRP doses when analyzed with automated squint, Mouse Grimace Scale, and orbital tightening (Fig 2B, middle and bottom rows, middle and right panels).

### Lack of photic blink or squint in C57BL/6J and CD1 mice

An unexpected observation during these studies was that both C57BL/6J and CD1 mice appeared to lack a photic blink in response to bright light stimulation. We confirmed the lack of photic blink with the automated squint assay. Automated squint measured every 0.1 seconds is shown for the dark period (0.5 seconds prior to light exposure) and the light period (initial light stimulus to 0.5 seconds post-light exposure) in C57BL/6J mice (Fig. 3A, left panel) and CD1 mice (Fig. 3B, left panel). Automated squint for both strains shows no significant differences between pre and post-light stimulus (Fig. 3A, B right panels).

**Figure 3:**
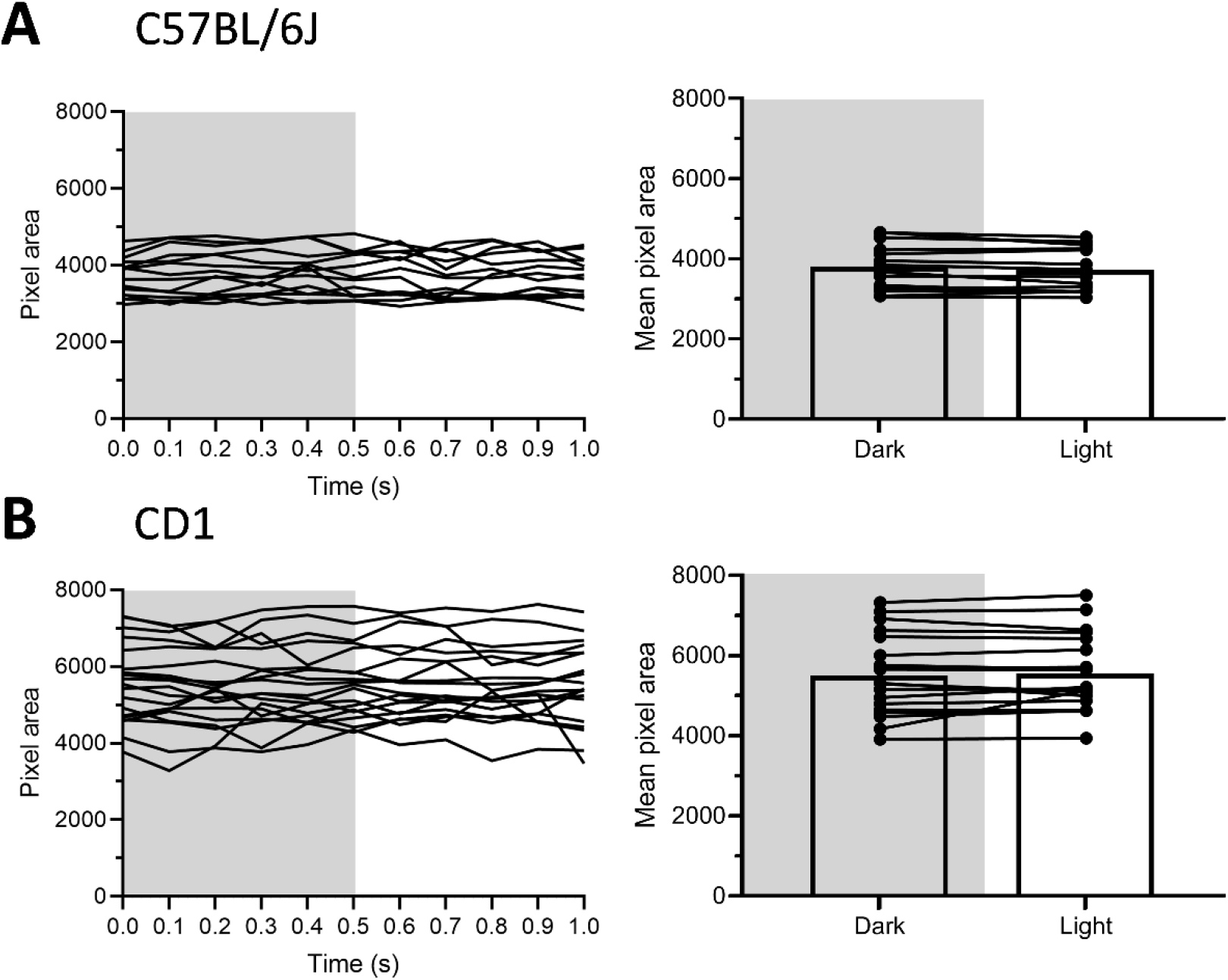
Automated squint analysis showing lack of photic blink in C57BL/6J and CD1 mice. Mice were restrained and recorded in a dark condition (∼0 lux) then exposed to a continuous white light exposure (1250 lux, 5600k). (**A**) Right eye palpebral fissure pixel area for individual C57BL/6J (black) mice (n = 14) in a dark condition (shaded region) followed by a white light exposure. The left and right panels show the individual tracings over time and the mean pixel area for each mouse at each condition, respectively. (**B**) Right eye palpebral fissure pixel area for individual CD1 (white) mice (n = 19) in a dark condition (shaded region) followed by a white light exposure. The left and right panels show the individual tracings over time and the mean pixel area for each mouse at each condition, respectively. Paired t-test revealed no significant differences between dark and light stimulation (C57BL/6J mice: p = 0.069 n = 14, CD1 mice: p = 0.4347, n = 19).

To confirm the lack of photic blink in mice, we tested transgenic CGRP-sensitized nestin/hRAMP1 mice by measuring the EMG of the orbicularis oculi muscle. The nestin/hRAMP1 mice were chosen to increase the likelihood of detecting a light-induced response based on their increased CGRP-induced light sensitivity in the light aversion assay [17]. A control littermate mouse displayed no EMG response when transitioning from dark to light stimulus (745 lux, 455 nm) with the same recording duration as the automated squint assay (Fig 4A, left panel). As a positive control, the same control littermate mouse did show an EMG response to an air puff (Fig 4A, right panel, black arrow) post-light stimulus which caused a blink. Nestin/hRAMP1 double transgenic mice displayed the same behavior as the control littermate (Fig. 4B) demonstrating these mice lack a photic blink yet display a reflexive blink in response to a corneal air puff stimulus that can be detected through EMG.

**Figure 4:**
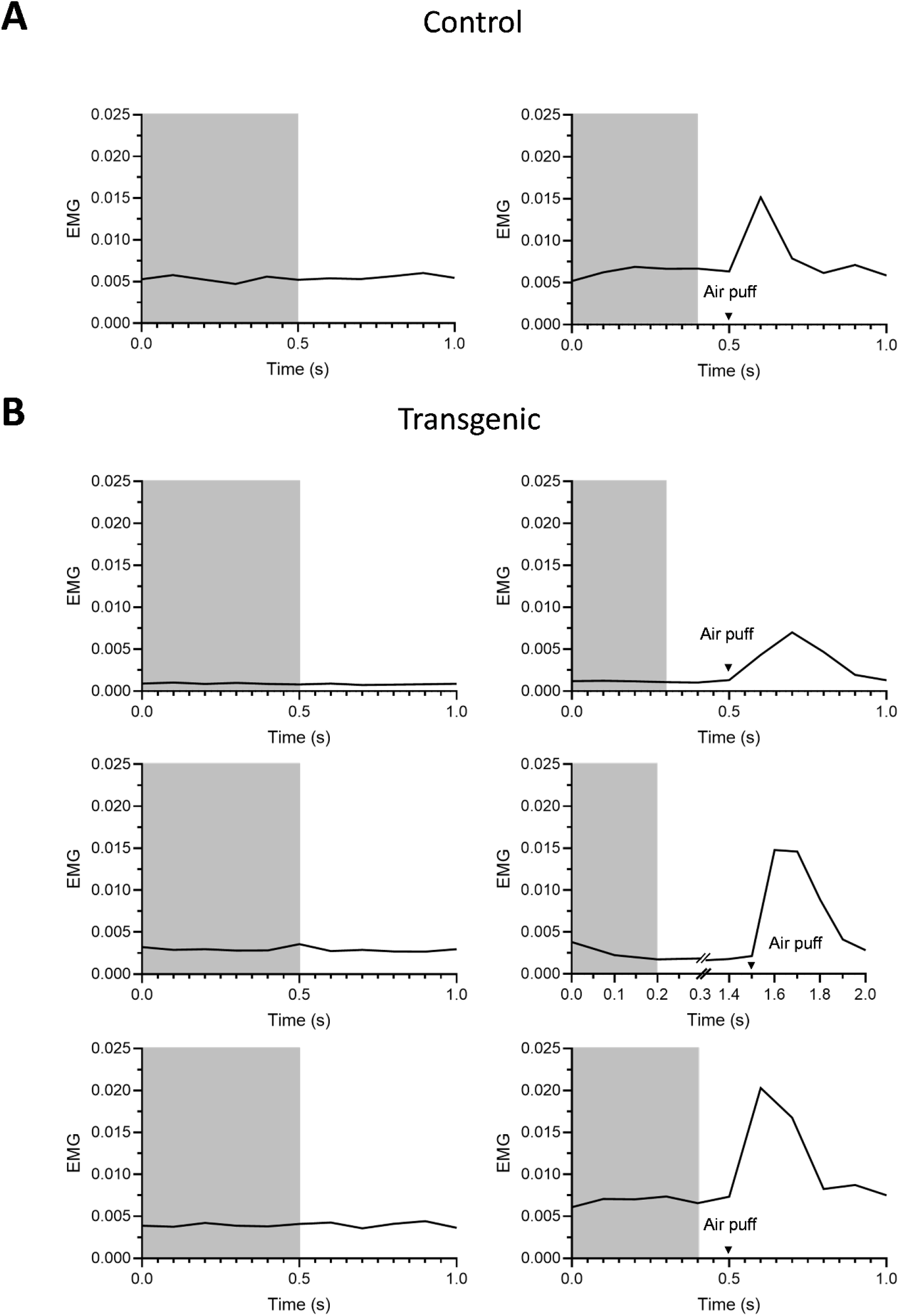
EMG activity of the orbicularis oculi muscle in response to light and air puff in nestin/hRAMP1 and littermate mice. (**A**) One control littermate was subcutaneously implanted with a radio telemeter with leads inserted into the orbicularis oculi muscle of the left eye. After 24 hours post-operation, mice were recorded during a baseline dark condition (∼0 lux shaded region) followed by a light stimulus (745 lux, 455 nm, unshaded region) without ipsilateral corneal air puff (left panel). Following reacclimation to the dark (∼0 lux) for 15 seconds, a second light stimulus was administered along with an ipsilateral corneal air puff (right panel, black arrow). Resulting EMG activity is presented as RMS in 100 ms windows (every 0.1 seconds). The corneal air puff caused a spike in EMG activity. (**B**) Three nestin/hRAMP1 double transgenic mice were analyzed in the same manner as the control littermate and displayed similar recordings.

### Automated squint detection of spontaneous pain induced by the peptide hormone amylin

With the validation of the automated squint assay, we measured the response of C57BL/6J mice to amylin, a peptide hormone closely related to CGRP. Human amylin has ∼50% sequence identity with CGRP and the two peptides bind a shared receptor (AMY1) with equal affinity [2]. Mean pixel area over time for baselines and respective treatments as well as AUC are shown in Fig 5A. C57BL/6J mice displayed significant differences in squint with 0.5 mg/kg amylin compared to baseline, while vehicle-treated mice showed no difference in squint response compared to respective baselines (Fig 5B). When viewed separately, a sex difference was detected with amylin-induced squint response. The female C57BL/6J mice displayed a significant squint response when compared to respective baselines while the males did not have a detectable response (Fig. 5C).

**Figure 5:**
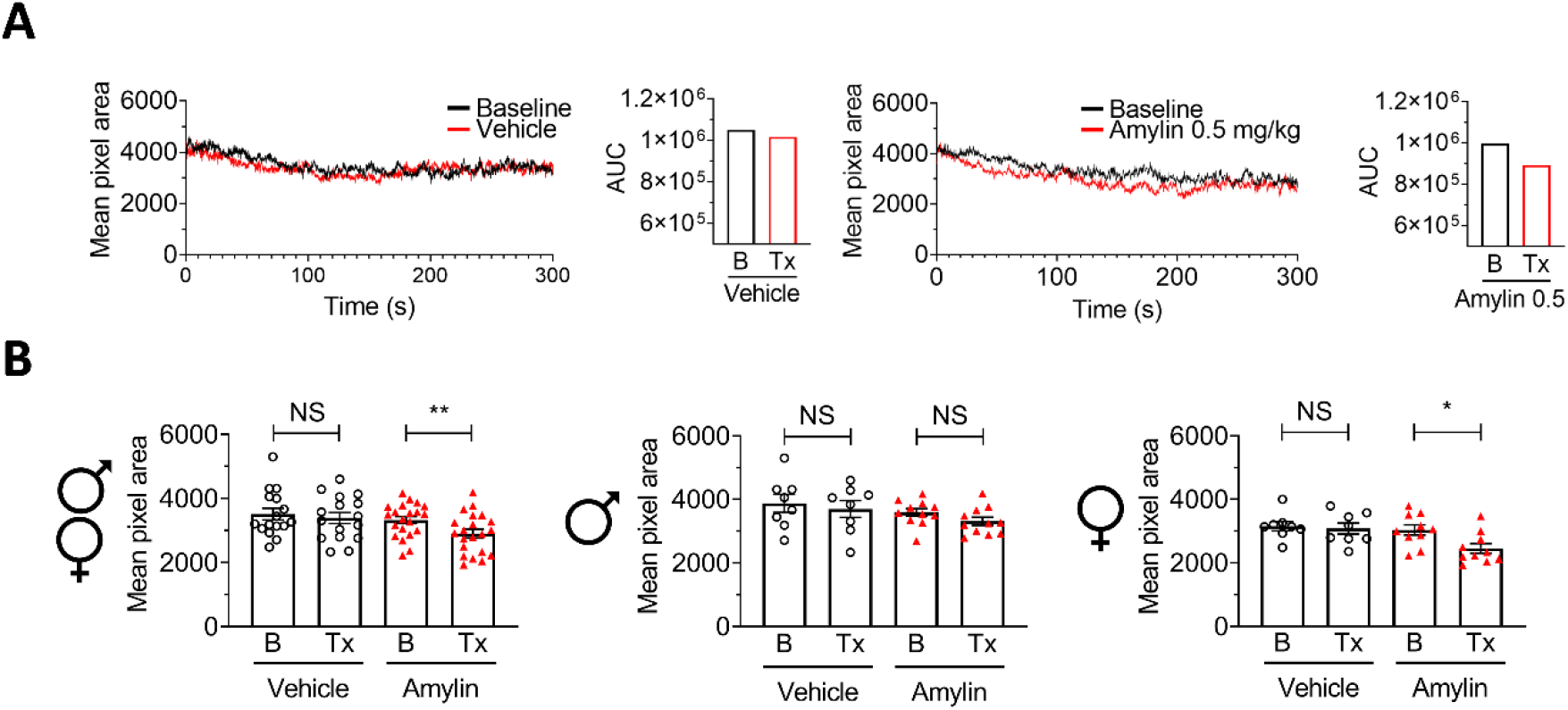
Amylin induces facial grimace in female but not male C57BL/6J mice. (**A**) Mean pixel area over time for either vehicle (n = 16, PBS, left panel) or amylin (n = 21, 0.5 mg/kg, right panel). Area under the curve for the averages are shown to the right of the line graphs. (**B**) Mean overall pixel area scores from panel A for all mice during baseline (B) and treatment (Tx) conditions. The data are shown for all mice (left panel), male only (middle panel, n = 8, PBS; n = 11, amylin), and female only (right panel, n = 8, PBS; n = 10 amylin). Two way repeated-measures ANOVA followed by Šídák’s multiple-comparison test to compare baseline and treatment conditions, ^*^p < 0.05, ^**^p < 0.01.

## DISCUSSION

In this report, we describe an automated system for measuring squint as a faster, more efficient readout of facial pain in mice in response to CGRP. The software was able detect and measure squint activity in both black C57BL/6J mice and white CD1 mice, demonstrating a range of application for squint analysis. We validated its accuracy by comparing automated and manual scoring of CGRP-induced grimace in C57BL/6J mice, showing a greater sensitivity of the automated squint response to low dose CGRP, with a response detected in female mice but not males. In addition, we used the automated squint assay to show that mice lack a photic blink response which was confirmed with EMG recordings of orbicularis oculi muscle activity. Finally, we used the automated assay to document that amylin induces a small but significant squint response in female C57BL/6J mice. These results demonstrate the applicability of a new automated method for detection of facial changes and specifically squinting associated with pain.

The ability of amylin to induce squint in mice builds on the recent report that the amylin analog pramlintide induces migraine-like headache in patients and that amylin induced two migraine-like symptoms of light aversion and cutaneous hypersensitivity in mice [5]. While our experiments were not designed for direct comparison between CGRP and amylin, the relatively small amylin-induced squint response is consistent with the observations that pramlintide and amylin also appeared to be less potent than CGRP in inducing migraine-like headaches and symptoms in patients and mice, respectively [5]. Future studies are needed to ascertain whether the difference in efficacy might reflect differences in receptors i.e., CGRP binds the canonical CGRP receptor to a greater degree than amylin [8].

Our results highlight the use of the eye as an accurate measurement of spontaneous pain in a CGRP or amylin-evoked model. When compared directly to the facial grimace score, the automated squint detection was able to identify smaller changes than the human scored Mouse Grimace Assay. Of importance, the automated analysis enabled the detection of a sex difference, with females showing a response to low dose CGRP in our mouse model that was not detected manually with the Mouse Grimace Scale. This is consistent with our previous study that found orbital tightening to be the principal component in mouse facial grimace [16].

The development of automated pain analysis software is quickly evolving. Tuttle et al trained a convolutional neural network with mouse images previously evaluated using the Mouse Grimace Scale to distinguish the presence of pain/no-pain in a binary assessment [20]. However, small changes in pain state on a continuous scale were not possible. When using convolutional neural networks, the differential characteristics detected by the software are unbeknownst to the user. The automated system reported here utilized a definable unit (palpebral fissure area) on a continuous scale that was statistically supported as a read out for CGRP or amylin-induced pain. As more video frames of faces are landmarked and incorporated into the feature detection model, the automated system described here will be further optimized in its ability to detect pain states in mice utilizing more facial features in addition to squint. Additionally, this method and software can be applied to other species, including humans, in different environments and with different video recording systems. Detection of squint that can be objectively quantified in real time with low computing power as a read-out of pain is a key difference from neural network-based systems used to detect changes in facial features.

Other machine learning paradigms are being used to detect sub-millisecond changes in behavior, which will likely lead to a wealth of information. Indeed, automated tracking of evoked pain responses such as with foot touch stimulation will nicely pair with the automated squint assay. The tracked evoked foot response images published by Jones et al also showed varying degrees of squinting over the range of stimuli [9]. Thanks to high frame rate recording, these machine learning paradigms can detect time-resolved information undetectable by human observer-based assays. Automated methods to encompass pain detection by facial feature analysis will provide more sensitive and translatable readouts of pain analysis that could greatly increase the probability of drug discovery [15] applicable to human and veterinary medicine.

While advances have been made in this automated model, hurdles to overcome exist within the system. A major caveat is that this version of the automated squint software requires the mouse to be in a fixed position from the array of video cameras in order to keep the magnification constant between mice and compared to baseline. Restraint is a known inducer of stress in animals and may lead to a blunting of effects, as noted in our previous study [16]. Future efforts will focus on quantifying facial features in unrestrained mice. Direct measurement of the millimeter distance between the inner and outer canthus of the palpebral fissure of each animal will allow proper magnification scaling in free roaming mice so that restraint will not be a requirement.

Development of the automated squint analysis did allow us to make two unexpected findings. The first was detection of a sex difference at a low dose of CGRP that we have not previously observed with the manual grimace assay [16], or with cutaneous sensitivity or light aversion assays [13; 17; 21]. In addition, amylin was effective at inducing squint in only female mice, even at the relatively high single dose that was tested. A similar observation of greater responses in female mice to the same dose of amylin has been reported with light aversion and cutaneous sensitivity tests [5]. While the reason for the greater sensitivity to IP CGRP and amylin in female mice is not known, it is interesting that Dussor and colleagues reported that only female mice responded to direct application of CGRP onto the dura [1]. This raises the possibility that perhaps at low CGRP doses, the site of action is preferentially at the dura, while other sites of action are recruited at higher doses. By extension, this suggests that amylin may also preferentially work in the dura. Future studies are needed to test these hypotheses.

The second unexpected finding was the lack of a photic blink reflex in mice. This is in contrast to the robust photic blink response in humans [7; 18; 23]. To our knowledge the ability of a light stimulus to trigger a blink in mice or rats has not been previously tested. However, light modulation of air puff-evoked blinks and EMG activity has been used in a photophobia study by Evinger and colleagues, who observed that bright light increased the spontaneous blink rate and magnitude of air puff-evoked blinks in rats [4]. Along a similar line, glycerol trinitrate, a trigger of migraine-like symptoms in rodents, potentiated air puff-evoked blink-associated EMG activity in anesthetized rats [10]. So, while the blink reflex can be modulated by light and a migraine trigger in rats, at least in mice we find that light itself is not sufficient to trigger a squint or EMG response. Furthermore, we previously found that light did not affect the squint response to CGRP [16].

In summary, this study is a step forward in the development of a quantifiable translational pain assay. The high throughput nature of the automated squint assay greatly reduces turnaround time for analysis after data collection and provides a continuous scale readout of a measurable component of the grimace response associated with pain. The applicability of the assay is shown by the more sensitive detection of squint responses to CGRP and the ability to detect responses to amylin in female mice.

## References

[1] Avona A, Burgos-Vega C, Burton MD, Akopian AN, Price TJ, Dussor G. Dural Calcitonin Gene-Related Peptide Produces Female-Specific Responses in Rodent Migraine Models. J Neurosci 2019;39(22):4323–4331.

[2] Bower RL, Hay DL. Amylin structure-function relationships and receptor pharmacology: implications for amylin mimetic drug development. Br J Pharmacol 2016;173(12):1883–1898.

[3] Crosta QR, Ward TM, Walker AJ, Peters LM. A review of pain measures for hospitalized children with cognitive impairment. J Spec Pediatr Nurs 2014;19(2):109–118.

[4] Dolgonos S, Ayyala H, Evinger C. Light-induced trigeminal sensitization without central visual pathways: another mechanism for photophobia. Invest Ophthalmol Vis Sci 2011;52(11):7852–7858.

[5] Ghanizada H, Al-Karagholi MA, Walker CS, Arngrim N, Rees T, Petersen J, Siow A, Morch-Rasmussen M, Tan S, O’Carroll SJ, Harris P, Skovgaard LT, Jorgensen NR, Brimble M, Waite JS, Rea BJ, Sowers LP, Russo AF, Hay DL, Ashina M. Amylin analog pramlintide induces migraine-like attacks in patients. Ann Neurol 2021.

[6] Glaros AG, Marszalek JM, Williams KB. Longitudinal Multilevel Modeling of Facial Pain, Muscle Tension, and Stress. J Dent Res 2016;95(4):416–422.

[7] Hackley SA, Johnson LN. Distinct early and late subcomponents of the photic blink reflex: response characteristics in patients with retrogeniculate lesions. Psychophysiology 1996;33(3):239–251.

[8] Hay DL, Garelja ML, Poyner DR, Walker CS. Update on the pharmacology of calcitonin/CGRP family of peptides: IUPHAR Review 25. Br J Pharmacol 2018;175(1):3–17.

[9] Jones JM, Foster W, Twomey CR, Burdge J, Ahmed OM, Pereira TD, Wojick JA, Corder G, Plotkin JB, Abdus-Saboor I. A machine-vision approach for automated pain measurement at millisecond timescales. Elife 2020;9.

[10] Jones MG, Andreou AP, McMahon SB, Spanswick D. Pharmacology of reflex blinks in the rat: a novel model for headache research. J Headache Pain 2016;17(1):96.

[11] Lalloo C, Stinson JN. Assessment and treatment of pain in children and adolescents. Best Pract Res Clin Rheumatol 2014;28(2):315–330.

[12] Langford DJ, Bailey AL, Chanda ML, Clarke SE, Drummond TE, Echols S, Glick S, Ingrao J, Klassen-Ross T, Lacroix-Fralish ML, Matsumiya L, Sorge RE, Sotocinal SG, Tabaka JM, Wong D, van den Maagdenberg AM, Ferrari MD, Craig KD, Mogil JS. Coding of facial expressions of pain in the laboratory mouse. Nat Methods 2010;7(6):447–449.

[13] Mason BN, Kaiser EA, Kuburas A, Loomis MM, Latham JA, Garcia-Martinez LF, Russo AF. Induction of Migraine-Like Photophobic Behavior in Mice by Both Peripheral and Central CGRP Mechanisms. J Neurosci 2017;37(1):204–216.

[14] Mogil JS, Pang DSJ, Silva Dutra GG, Chambers CT. The development and use of facial grimace scales for pain measurement in animals. Neurosci Biobehav Rev 2020;116:480–493.

[15] Pereira TD, Aldarondo DE, Willmore L, Kislin M, Wang SS, Murthy M, Shaevitz JW. Fast animal pose estimation using deep neural networks. Nat Methods 2019;16(1):117–125.

[16] Rea BJ, Wattiez AS, Waite JS, Castonguay WC, Schmidt CM, Fairbanks AM, Robertson BR, Brown CJ, Mason BN, Moldovan-Loomis MC, Garcia-Martinez LF, Poolman P, Ledolter J, Kardon RH, Sowers LP, Russo AF. Peripherally administered calcitonin gene-related peptide induces spontaneous pain in mice: implications for migraine. Pain 2018;159(11):2306–2317.

[17] Recober A, Kaiser EA, Kuburas A, Russo AF. Induction of multiple photophobic behaviors in a transgenic mouse sensitized to CGRP. Neuropharmacology 2010;58(1):156–165.

[18] Tavy DL, van Woerkom TC, Bots GT, Endtz LJ. Persistence of the blink reflex to sudden illumination in a comatose patient. A clinical and pathologic study. Arch Neurol 1984;41(3):323–324.

[19] Tracey I, Woolf CJ, Andrews NA. Composite Pain Biomarker Signatures for Objective Assessment and Effective Treatment. Neuron 2019;101(5):783–800.

[20] Tuttle AH, Molinaro MJ, Jethwa JF, Sotocinal SG, Prieto JC, Styner MA, Mogil JS, Zylka MJ. A deep neural network to assess spontaneous pain from mouse facial expressions. Mol Pain 2018;14:1744806918763658.

[21] Wattiez AS, Sowers LP, Russo AF. Calcitonin gene-related peptide (CGRP): role in migraine pathophysiology and therapeutic targeting. Expert Opin Ther Targets 2020;24(2):91–100.

[22] Williams AC. Facial expression of pain: an evolutionary account. Behav Brain Sci 2002;25(4):439-455; discussion 455-488.

[23] Yates SK, Brown WF. Light-stimulus-evoked blink reflex: methods, normal values, relation to other blink reflexes, and observations in multiple sclerosis. Neurology 1981;31(3):272–281.

[24] Zhang Z, Winborn CS, Marquez de Prado B, Russo AF. Sensitization of calcitonin gene-related peptide receptors by receptor activity-modifying protein-1 in the trigeminal ganglion. J Neurosci 2007;27(10):2693–2703.

